# Application of an Interpretable Classification Model on Early Folding Residues during Protein Folding

**DOI:** 10.1101/381483

**Authors:** Sebastian Bittrich, Marika Kaden, Christoph Leberecht, Florian Kaiser, Thomas Villmann, Dirk Labudde

## Abstract

**Background:** Machine learning strategies are prominent tools for data analysis. Especially in life sciences, they have become increasingly important to handle the growing datasets collected by the scientific community. Meanwhile, algorithms improve in performance, but also gain complexity, and tend to neglect interpretability and comprehensiveness of the resulting models.

**Results:** Generalized Matrix Learning Vector Quantization (GMLVQ) is a supervised, prototype-based machine learning method and provides comprehensive visualization capabilities not present in other classifiers which allow for a fine-grained interpretation of the data. In contrast to commonly used machine learning strategies, GMLVQ is well-suited for imbalanced classification problems which are frequent in life sciences. We present a Weka plug-in implementing GMLVQ. The feasibility of GMLVQ is demonstrated on a dataset of Early Folding Residues (EFR) that have been shown to initiate and guide the protein folding process. Using 27 features, an area under the receiver operating characteristic of 76.6% was achieved which is comparable to other state-of-the-art classifiers.

**Conclusions:** The application on EFR prediction demonstrates how an easy interpretation of classification models can promote the comprehension of biological mechanisms. The results shed light on the special features of EFR which were reported as most influential for the classification: EFR are embedded in ordered secondary structure elements and they participate in networks of hydrophobic residues. Visualization capabilities of GMLVQ are presented as we demonstrate how to interpret the results.

## Introduction

The analysis of data collected during biological experiments poses a challenge for modern bioinformatics. Usually this data is feature rich, yet hard to interpret, such as it is the case for single-cell gene expression data obtained by high-throughput experiments [1]. Despite sophisticated pre-processing and the application of machine learning models, analysis – and most importantly interpretation – of such data is still hard to accomplish. Nevertheless, machine learning is the basis for sophisticated predictions and allows new insights into open questions. In this paper, we examine the problem of protein folding, by the means of Early Folding Residues (EFR).

Further, we apply an interpretable classifier on this problem to deepen the understanding of EFR based on the trained model.

### Grasping the protein folding problem through Early Folding Residues

Proteins are chains of amino acids which are connected by covalent bonds and, for the most part, autonomously fold into a defined structure (Figure 1) [2, 3]. This stable, three-dimensional structure allows proteins to be functional and catalyze particular chemical reactions, transport molecules, or transduce signals in cells. The fundamentals of the so-called protein folding process are still unclear.

**Figure 1.**
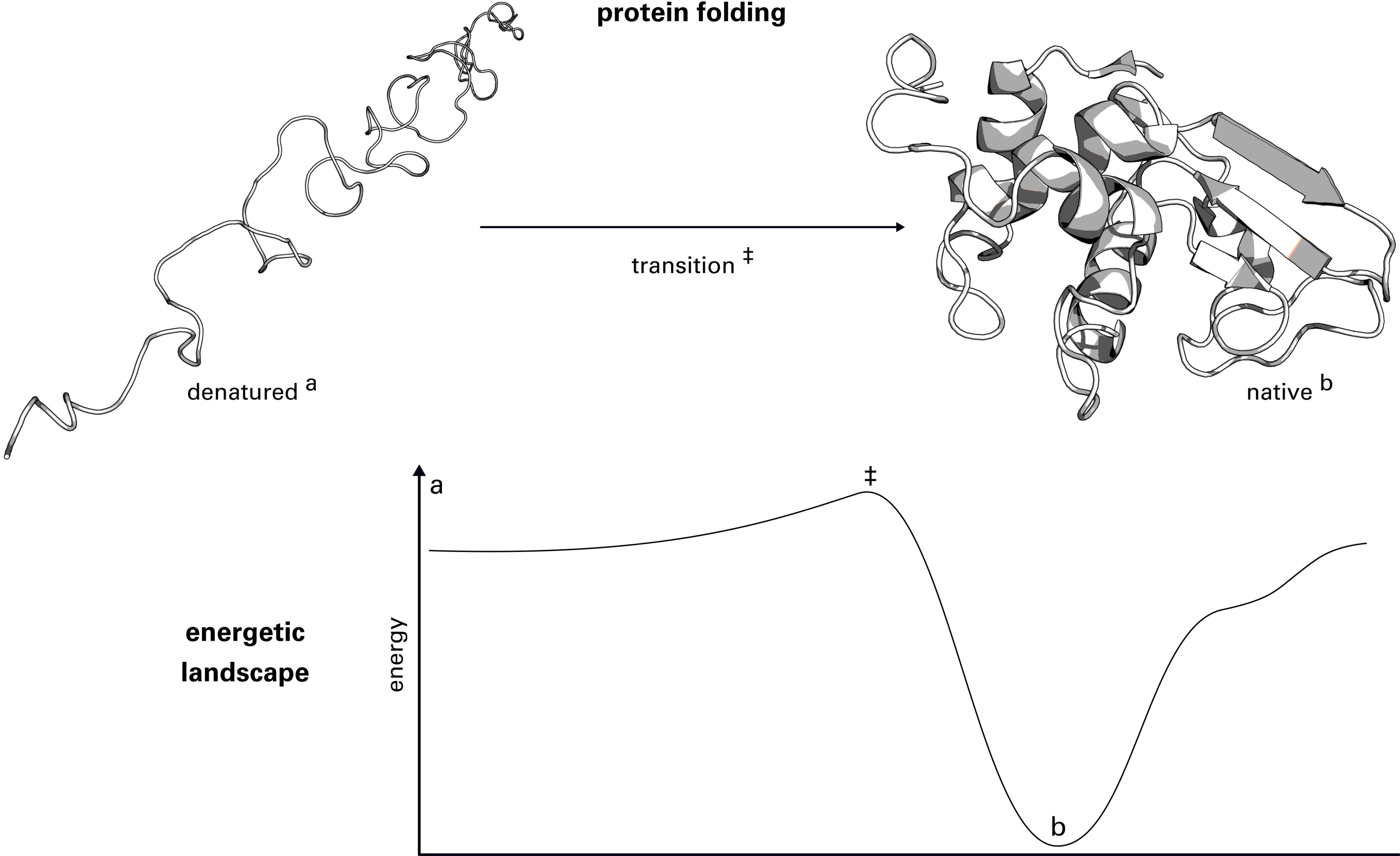
Illustration of the protein folding process. The denatured protein (a) has to pass an energetic barrier (‡), the so-called transition state, to reach its native three-dimensional structure (b). Usually, the native structure represents the global energetic optimum of the protein. EFR are residues which initiate and guide the folding process [13, 19].

Folding intermediates are highly unstable and thus protein folding was difficult to investigate experimentally for a long time [4, 5]. Nowadays, pulse labeling hydrogen-deuterium exchange is a prominent tool to investigate the folding process with spatial and temporal resolution [6–13]. EFR were identified as key residues for the folding process as they participate in the earliest folding events. By forming long-range tertiary contacts, EFR are also assumed to guide the assembly of different protein regions which stabilize the native, folded protein structure [10–13]. EFR were shown to be the initiators of the folding process and, thus, may be used to advance the understanding of the protein folding process [14–17]. The process remains elusive, but understanding the peculiarities of the folding nucleus [12, 16–19] (as indicated by EFR) may aid unraveling it by providing information on intermediate states of the process. The currently identified set of EFR is deposited in the Start2Fold database [13] which provides a robust dataset for the characterization of EFR and the design of classifiers for their prediction. Raimondi et al. developed the predictor *EFoldMine* that discriminates EFR from other residues, termed Late Folding Residues (LFR), using features derived from the protein sequence [16]. The aim of their paper was to distinguish between these two classes using secondary structure propensities and backbone rigidity values of surrounding sequence fragments. It is crucial to understand, which features cause a small number of residues to become EFR while the majority of them are LFR. Unfortunately, the classifiers applied by Raimondi et al. [16] cannot provide detailed insights and the published model is not discussed under this focus. This is mainly the consequence of the chosen features and the employed standard Support Vector Machine (SVM) with the Radial Basis Function (RBF)-kernel; this results in a model which is difficult to interpret and does not state the features relevant to distinguish the classes.

We created a dataset using the same data basis but utilize a more diverse set of features. This set includes information derived from the protein structure as a corresponding structure is deposited in the Protein Data Bank (PDB) for each protein of the dataset. This allows for a better interpretability and discussion of the resulting model and, thus, emphasizes unique aspects of the Generalized Matrix Learning Vector Quantization (GMLVQ) classifier. Our study demonstrates how an adaptation of an established machine learning strategy allows pinpointing the most influential features for classification. Therefore, we present a novel implementation of the GMLVQ algorithm [20, 21] as plug-in for the popular Waikato Environment For Knowledge Analysis (Weka) framework [22–24]. This plug-in features diverse visualization tools which encourage the user to interpret the resulting model and render GMLVQ a comprehensible *white box* classifier.

## Detailed description of the dataset of Early Folding Residues

The Start2Fold database [13, 19] contains the results of pulse labeling hydrogen-deuterium exchange experiments for 30 proteins. Due to the nature of the experimental data, no information can be obtained for the amino acid proline because its amide group is not susceptible to an exchange of hydrogen to deuterium. All 111 proline instances were dropped from the initial dataset which resulted in 3, 266 residues of which 482 (14.8%) are EFR.

### Feature annotation

We opted to describe every amino acid in the dataset by a number of features capturing different aspects of their molecular surroundings and physicochemical properties. Amino acids have sequential and spatial neighbors and both levels of organization are strongly intertwined by the process of protein folding [25]. All considered features describe a particularized aspect of this connection and are summarized in Table 1. Features of each residue were averaged with respect to four adjacent positions at the sequence level in N-as well as C-terminal region.

**Table 1.**
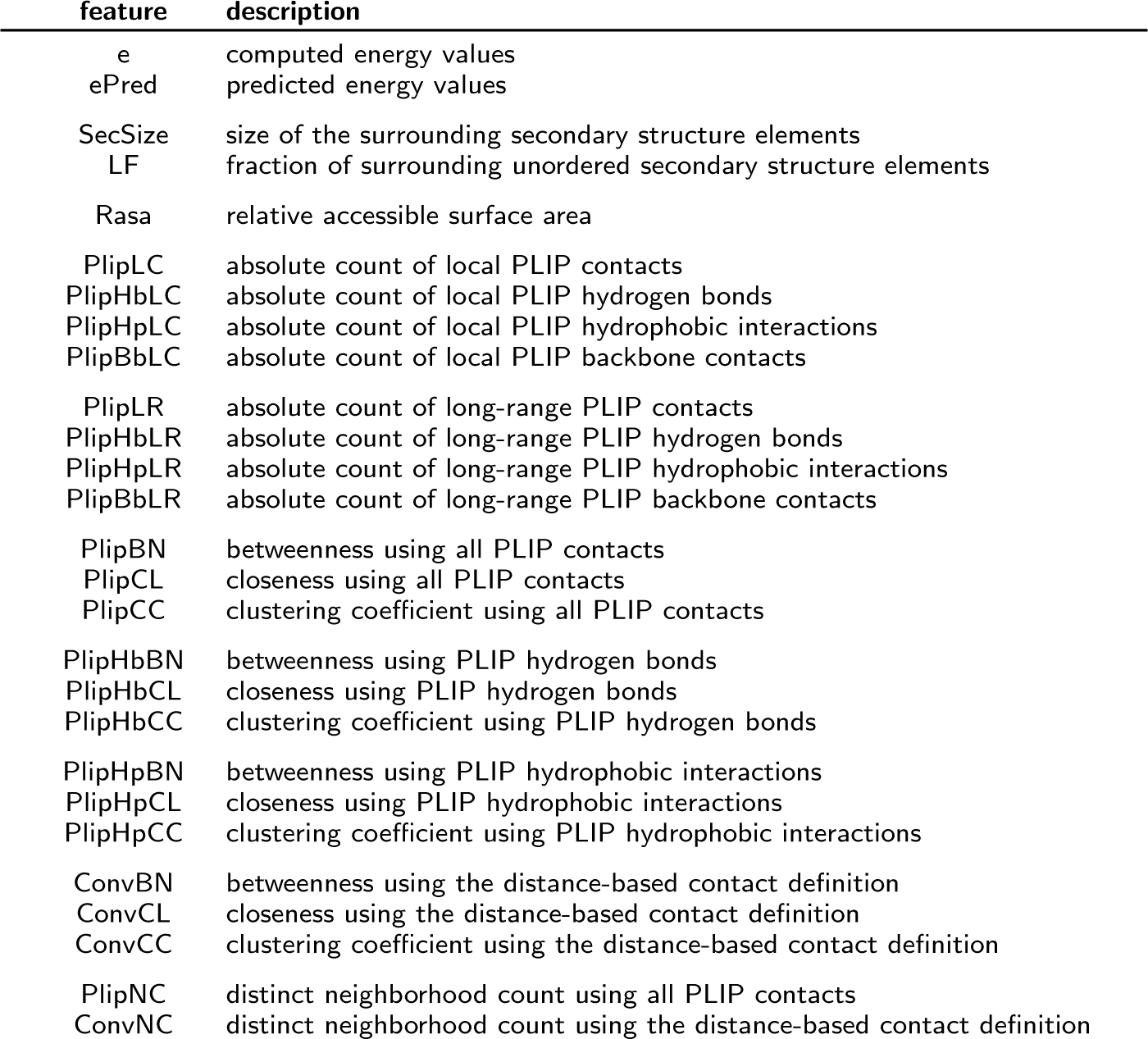
Denomination and short description of the 27 features of the dataset for individual residues classification. References to these features are given in *italic* font.

| feature | description |
| --- | --- |
| e | computed energy values |
| ePred | predicted energy values |
| SecSize | size of the surrounding secondary structure elements |
| LF | fraction of surrounding unordered secondary structure elements |
| Rasa | relative accessible surface area |
| PlipLC | absolute count of local PLIP contacts |
| PlipHbLC | absolute count of local PLIP hydrogen bonds |
| PlipHpLC | absolute count of local PLIP hydrophobic interactions |
| PlipBbLC | absolute count of local PLIP backbone contacts |
| PlipLR | absolute count of long-range PLIP contacts |
| PlipHbLR | absolute count of long-range PLIP hydrogen bonds |
| PlipHpLR | absolute count of long-range PLIP hydrophobic interactions |
| PlipBbLR | absolute count of long-range PLIP backbone contacts |
| PlipBN | betweenness using all PLIP contacts |
| PlipCL | closeness using all PLIP contacts |
| PlipCC | clustering coefficient using all PLIP contacts |
| PlipHbBN | betweenness using PLIP hydrogen bonds |
| PlipHbCL | closeness using PLIP hydrogen bonds |
| PlipHbCC | clustering coefficient using PLIP hydrogen bonds |
| PlipHpBN | betweenness using PLIP hydrophobic interactions |
| PlipHpCL | closeness using PLIP hydrophobic interactions |
| PlipHpCC | clustering coefficient using PLIP hydrophobic interactions |
| ConvBN | betweenness using the distance-based contact definition |
| ConvCL | closeness using the distance-based contact definition |
| ConvCC | clustering coefficient using the distance-based contact definition |
| PlipNC | distinct neighborhood count using all PLIP contacts |
| ConvNC | distinct neighborhood count using the distance-based contact definition |

#### Energy profiling

Energy Profiles [26, 27] transform the three-dimensional arrangement of atoms in a protein into a vector of energy values describing each amino acid. The computed energy (*e*) of a residues describes its interactions with its surroundings. Energy Profiles can also be predicted using only sequence information [26] (*ePred*) which represents the sequence composition. Computed as well as predicted energy values have been used before for the description of the folding process [26] as well as protein structure quality assessment [27].

#### Secondary structure elements

Secondary structure elements were annotated using DSSP [28] in its BioJava [29, 30] implementation. The secondary structure element size of a residue (*SecSize*) refers to the number of sequence neighbors sharing the same secondary structure (i.e. *α*-helix, *β*-strand, and coil). For sequence windows of nine residues the number of unordered secondary structure elements was counted and normalized by the window size [31]. This yields a fraction (*LF*), where high values are tied to regions of high disorder, whereas amino acids embedded in α-helices or *β*-sheets result in scores close to 0.

#### Relative accessible surface area

The Relative Accessible Surface Area (RASA) of a residue describes how exposed it is towards to solvent. Residues in the hydrophobic core tend to be buried and exhibit no accessible surface area. RASA values (*Rasa*) were computed with the BioJava [29, 30] implementation of the algorithm by Shrake and Rupley [32].

#### Non-covalent contacts

Non-covalent contacts stabilize protein structures and are the driving force behind protein folding [25]. The Protein-Ligand Interaction Profiler (PLIP) [33] was used for the annotation of non-covalent contacts between residues in protein structures. PLIP supports different contact types such as salt bridges, *π*-stacking interactions, or *π*-cation interactions. For this study, only hydrogen bonds (*Hb*) and hydrophobic interactions (*Hp*) were considered explicitly. Other contact types were not observed for most of the rather small proteins in the dataset. Furthermore, local and long-range contacts [34] were distinguished. Local contacts (suffix *LC*) are defined as contacts between residues less than six sequence positions apart – their main contribution is stabilizing secondary structure elements. In contrast, long-range contacts (suffix *LR)* occur between residues more than five sequence positions apart and constitute stabilizing contacts between secondary structure elements which primarily manifest the three-dimensional arrangement of a protein. Backbone contacts (*Bb*) occur only between backbone atoms of the respective residues.

#### Residue graph representation of proteins

Proteins in the dataset were represented as residue graphs. Amino acids always constituted the nodes and contacts between residues were represented by edges. Covalently bound residues were considered to be in contact. All contacts annotated by PLIP were used to create the first graph representation (using the prefix *Plip*). Reduced representations were created by only considering hydrogen bonds (prefix *PlipHb*) respectively hydrophobic interactions (prefix *PlipHp*). The contacts detected by PLIP may ignore spatially close residues when they do not form any contacts according the underlying rule set. Therefore, an additional contact definition was employed (prefix *Conv*): two residues were considered to be in contact, if their *C_α_* atoms were at most 8 Å apart.

#### Topological descriptors

Based on the four graph representations (*Plip, PlipHb, PlipHp,* and *Conv*), topological descriptors of individual residues were computed. This allows to describe how residues are connected to other residues. Most of these properties are based on shortest paths observable in the graph. The betweenness centrality (*BN*) of a node is defined as the number of shortest paths passing through that particular node. The term is normalized by the number of node pairs 0.5 ⋅ *n ⋅* (*n –* 1) in the residue graph with *n* nodes [35, 36]. The closeness centrality (*CL*) of a node is defined the inverse of the average path length to any other node. The clustering coefficient describes the surroundings of individual nodes. All adjacent nodes are collected and the number of edges between these *n_k_* nodes is determined. The clustering coefficient (*CC*) of a node is defined as number of edges between its adjacent nodes, divided by the maximum number of edges which can theoretically connect these nodes which is 0.5⋅*n_k_*⋅(*n_k_* – 1). The distinct neighborhood count (*NC*) captures how many sequentially distant (long-range) protein regions are connected by a residue [17].

## Description of the Generalized Matrix Learning Vector Quantization classifier

The Generalized Learning Vector Quantization (GLVQ) is a powerful distance- and prototype-based classification method [20]. The idea is adapted from the unsupervised vector quantizations methods such as k-Means or the Self-Organizing Map (SOM) and an extension of the heuristic Learning Vector Quantization (LVQ) [37]. For each class at least one prototype is initialized and a function, which approximates the classification accuracy (Figure 2), is maximized during learning. The optimization is commonly done by Stochastic Gradient Ascent (SGA) and ends up in an intuitive adaption of the prototypes. Thereby, in each iteration, for one training data point v two prototypes are taken into account: the nearest prototype with the same label as the data point and the nearest prototype with a different label, noted as **w**^+^(v) and **w**^−^(v). The prototype **w**^+^(v) is attracted while **w**^−^(v) is repulsed. The strength of attraction and repulsion is obtained by the gradients of the cost function and the according learning rates. The trained model is a nearest neighbor classifier, i. e. an incoming data point is assigned to the same class as the nearest prototype. In general, the GLVQ is a sparse model with interpretative prototypes. The complexity of the model can be chosen by the user by specifying the number of prototypes per class. If only one prototype per class and the Euclidean distance is applied, GLVQ is a linear classifier. A more detailed description of the algorithm can be found in [38, 39], Figure 3 provides a graphical representation.

**Figure 2.**
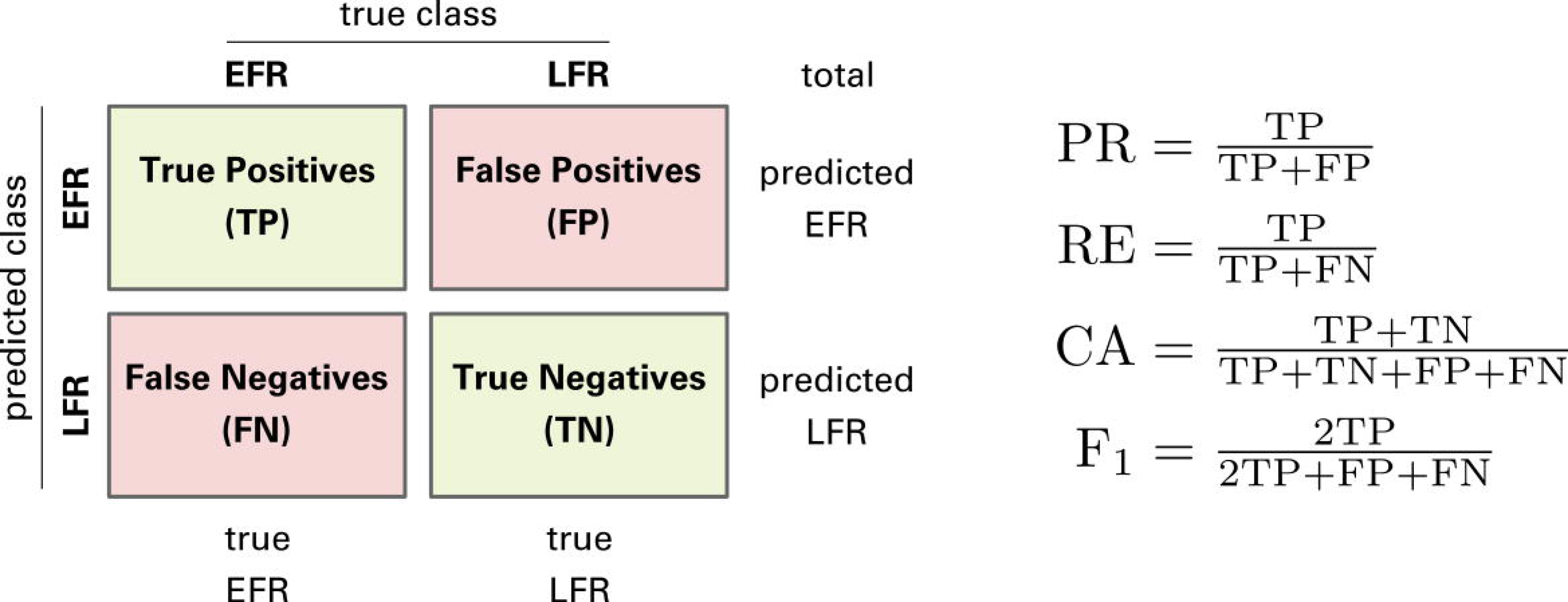
Confusion matrix. Exemplified Confusion Matrix with the formulas of precision (PR), recall (RE), accuracy (CA), and F_1_-measure.

**Figure 3.**
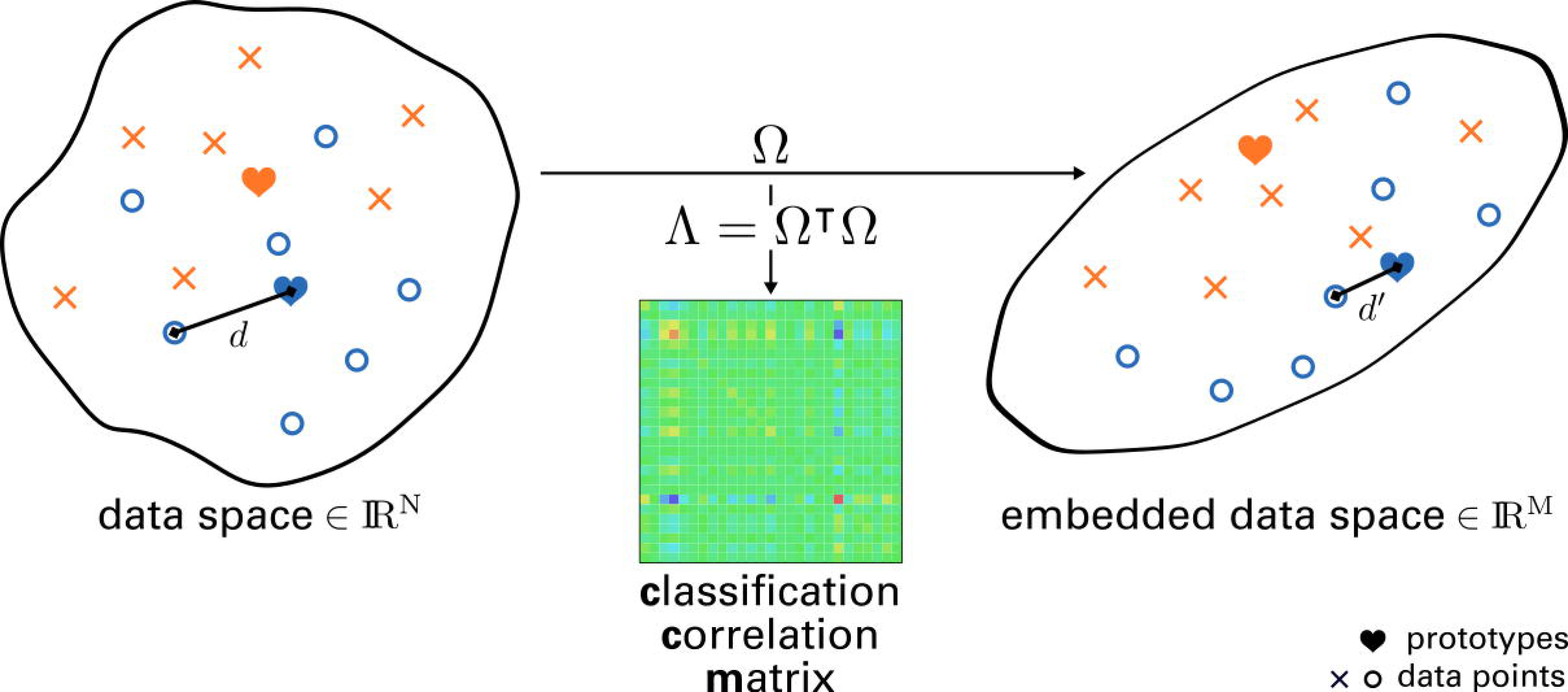
Principle of Generalized Matrix Learning Vector Quantization. Graphical depiction of learning with GMLVQ [51, 52]. One or multiple prototypes represent classes: each data point in the data space of dimension *N* belongs to the class of the prototype with the closest distance *d*. Prototypes are updated during learning as in LVQ [53]. Additionally, the matrix Ω maps the data space to an embedded data space of dimension *M*, where mapped distances *d*′ are optimized. The matrix Λ = Ω′Ω (CCM) represents the impact of each feature on the classification performance.

A prominent extension of the GLVQ is the Matrix GLVQ [21]. Beside the prototypes, a mapping of the data points is learned for better separation of the classes (Figure 4). This linear mapping, denoted by Ω ∈ ℝ^*m*×*D*^ is powerful and provides additional information about the classification problem. Thereby, D is the number of features. The parameter m can be chosen by the user and indicates the mapping dimension. If the mapping dimension is equal to D, the matrix is quadratic, but m can also be set to values smaller than *D*, e. g. down to *m* = 2. In the latter case the GMLVQ can be used for visualization of the dataset by mapping the dataset into the two-dimensional space [40]. Moreover, the matrix *CCM* = Ω*’*Ώ is termed Classification Correlation Matrix (CCM) [38]. In contrast to the correlation matrix of the features, the CCM reflects the correlations between the under the aspect of class discrimination (Figure 5B), i. e. positive or negative values of high magnitude between two features indicate a high positive or negative correlation of the features beneficial for the discrimination of classes. High values on the main diagonal occur for features important for the distinction of classes in general (see Figure 5A).

**Figure 4.**
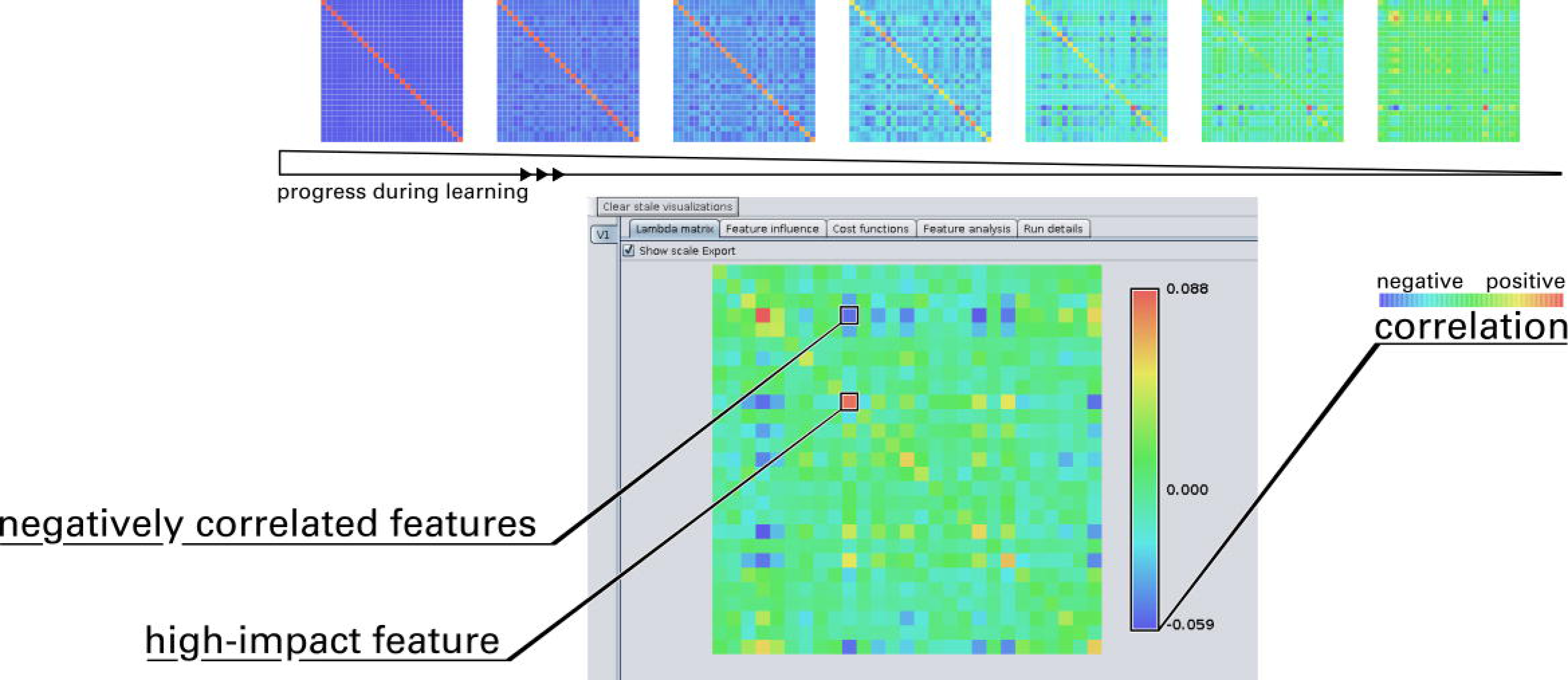
The process of learning. The graphical user interface of the GMLVQ Weka implementation. The matrix panel shows the CCM and displays live updates during the learning process. A color bar represents the scale of the matrix elements with a coloring scheme similar to a heat map.

**Figure 5.**
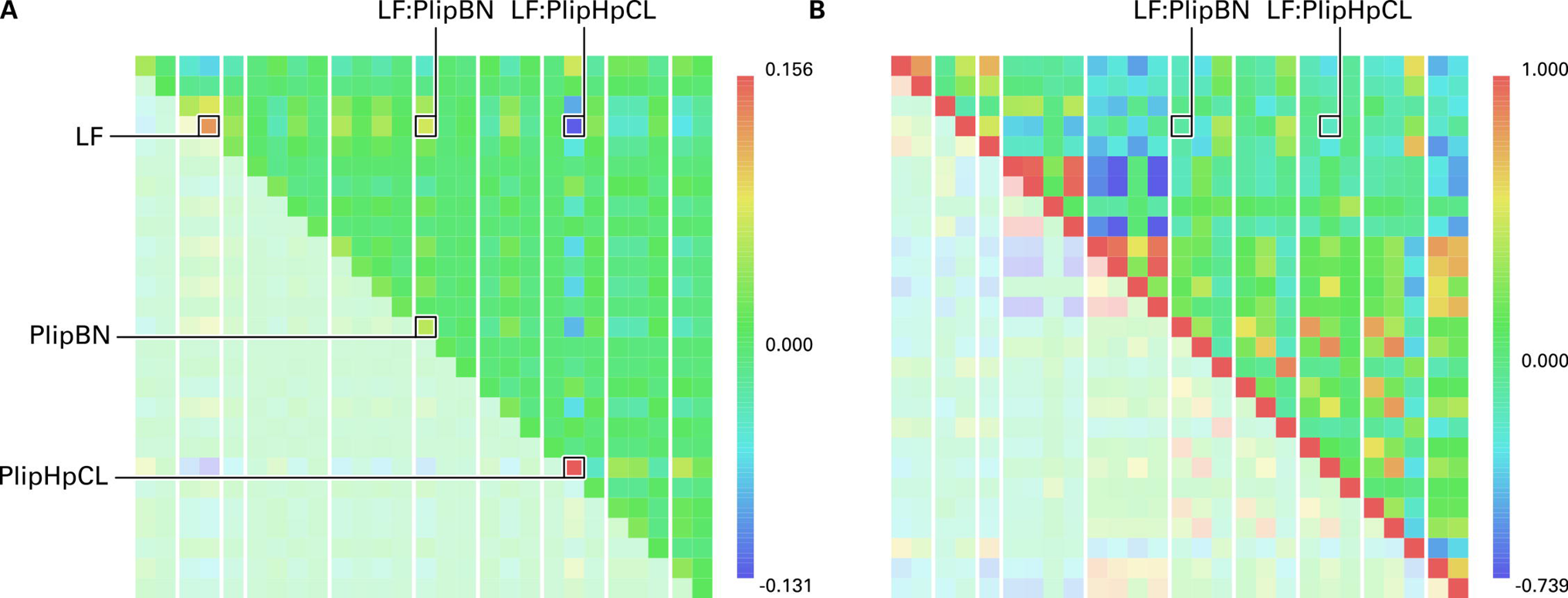
The Classification Correlation Matrix for the classification of Early Folding Residues. (A) The CCM depicts the positive impact of individual features for the classification performance on its main diagonal. Especially, PlipHpCL, LF, and PlipBN are features which discriminate EFR and LFR. The influence of ordered secondary structure elements was shown before [17, 19]. Both betweenness and closeness centrality tend to be increased for EFR which indicates their importance for the assembly of secondary structure elements by long-range hydrophobic interactions [17]. Other entries of the matrix describe pairs of features which are positively (red) or negatively (blue) correlated and increase classification performance further. (B) The standard correlation matrix of all features of the whole dataset. Again, positive and negative correlations are depicted in red and blue respectively. Interestingly, the features pointed out by GMLVQ do not stand out. Vice versa, strong correlations between features do not imply a favorable influence on the classification performance.

## Classification of Early Folding Residues

In the first step the dataset is standardized by z-score transformation. As mentioned before, the given dataset has a very unbalanced class distribution, i. e. only 482 data points of class EFR and 2784 of class LFR. In such cases the classification accuracy is inconclusive because it only takes correctly classified data points into account. Therefore, we determine further prominent evaluations measures based on the Confusion Matrix (CM) such as precision, recall, *F*_1_-measure, and Area Under The Receiver Operating Characteristic (auROC) [41, 42]. The precision considers data points predicted as the positive class (here EFR) and recall on all data points, which are real positives. In our example, the number of EFR is drastically smaller than that of LFR, so in general the precision is much worse than recall. The *F*_1_ - measure, which is the harmonic mean between precision and recall, is sensitive if one of these values is getting too small. The Receiver Operating Characteristic (ROC) is a graphical plot illustrating the trade-off between true positives and false positives for a model. According to the Weka documentation, the ROC is obtained by varying the threshold on the class probability estimates.

We applied 10-fold cross validation on different classifiers to compare the results of GMLVQ to other state-of-the-art methods (see Table 2). Furthermore, we investigate different parameter settings of the GMLVQ. On one side, the model size of the GMLVQ is a parameter chosen by the user. Here, we chose one prototype per class resulting in a linear classifier and five prototypes per class, which is more complex. Moreover, the GMLVQ has the feature to optimize other CM-based evaluation measures like the *F_β_*-measure or a linear combination of precision and recall. These can take the unbalanced class distribution into account. These aspects are reflected in Table 2. The comparison of the different classification models is challenging. It is difficult to decide objectively which classifier performs best. The SVM ends up with the best accuracy, yet the recall is very low. On the other side, the GMLVQ optimizing the weighted accuracy has the best recall and *F*_1_-value and optimizing the *F_β_*-measure ends up with the best value in the auROC. Furthermore, we can notice that very complex models do not automatically perform better. The Naive Bayes (NB), a very simple, fast and linear classifier performs comparable to the other much more complex models like Random Forest (RF) or SVM, which utilizes 1193 support vectors, i. e. 36% of the data points are necessary to describe the hyperplane. The GMLVQ runs with five prototype per class perform better in training than GMLVQ with one prototype, yet, in test the sparse model is more suitable.

**Table 2.**
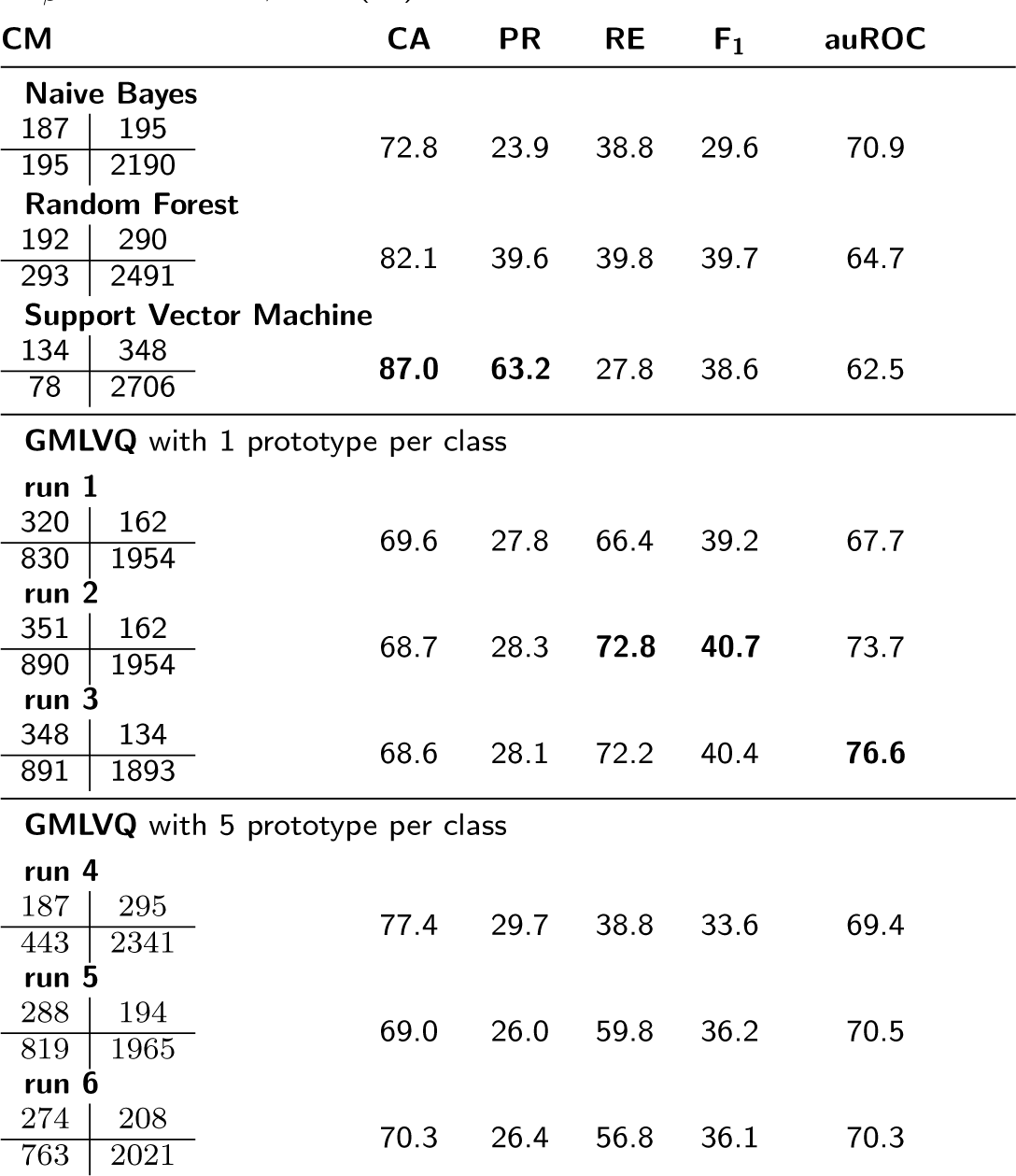
The test results in % right of confusion matrix (CM) and algorithmic parameters used for the classification of the data determined with Weka. Additionally, we marked the best values for the single evaluation measured bold. If not stated otherwise, default setup was used. SVM with RBF-kernel (*σ* = 5) which results in 1193 number of support vectors. Weights for weighted accuracy: 0.75 and 0.25. *F_β_*-measure with *β* = 1 (F_1_).

We applied different cost functions evaluating approximated values of classification accuracy, weighted classification accuracy, F_1_-measure, or weighted precision-recall. The results with the according parameter selection (Table 3) are listed in Table 2.

**Table 3.** Parameter selection to obtain the results of Table 2 using the Weka plug-in. Classification accuracy (CA), weighted classification accuracy (WCA) with weights 0.75 and 0.25 as well as F_*β*_-measure with *β* = 1 (F_1_).

| parameter | run 1 | run 2 | run 3 | run 4 | run 5 | run 6 |
| --- | --- | --- | --- | --- | --- | --- |
| cost function to optimize | CA | WCA | $F_1$ | CA | WCA | $F_1$ |
| number of epochs | 150 | 150 | 150 | 250 | 250 | 250 |
| number of prototypes | 1 | 1 | 1 | 5 | 5 | 5 |
| data point ratio per round | 0.75 | 0.75 | 0.75 | 0.75 | 0.75 | 0.75 |
| sigmoid sigma interval | [1.0,5.0] | [1.0,15.0] | [1.0,50.0] | [1.0,5.0] | [1.0,15.0] | [1.0,50.0] |
| prototype learning rate | 1.0 | 1.0 | 1.0 | 1.0 | 1.0 | 1.0 |
| matrix learning | true | true | true | true | true | true |
| omega learning rate | 1.0 | 1.0 | 1.0 | 1.0 | 1.0 | 1.0 |
| omega dimension | 27 | 27 | 27 | 27 | 27 | 27 |
| cost function beta | - | - | 1 | - | - | 1 |
| cost function weights | - | [0.75,0.25] | - | - | [0.75,0.25] | - |
| parallel execution | true | true | true | true | true | true |

To sum up, GMLVQ provides better results in recall even if the model is chosen to be very sparse. Distinguishing EFR and LFR is challenging and a clear separation was not achievable using the described features. GMLVQ was trained on the dataset in order to retrieve the most discriminative features of EFR and to showcase the capabilities and handling of the visualization.

### Visualization of learning process and interpretation of classification results

The GMLVQ plug-in tracks and summarizes each run by various visualization panels (Figure 6): the CCM panel (Figure 6A), the cost function panel (Figure 6B), the feature influence panel (Figure 6C), the feature analysis panel which depicts the prototype placement (Figure 6D), and the run details panel which reports the parameters of the corresponding run (Figure 6E). A detailed description on the example for the EFR dataset is given in order to demonstrate how results of GMLVQ can be interpreted by integrating information of these visualization panels.

**Figure 6.**
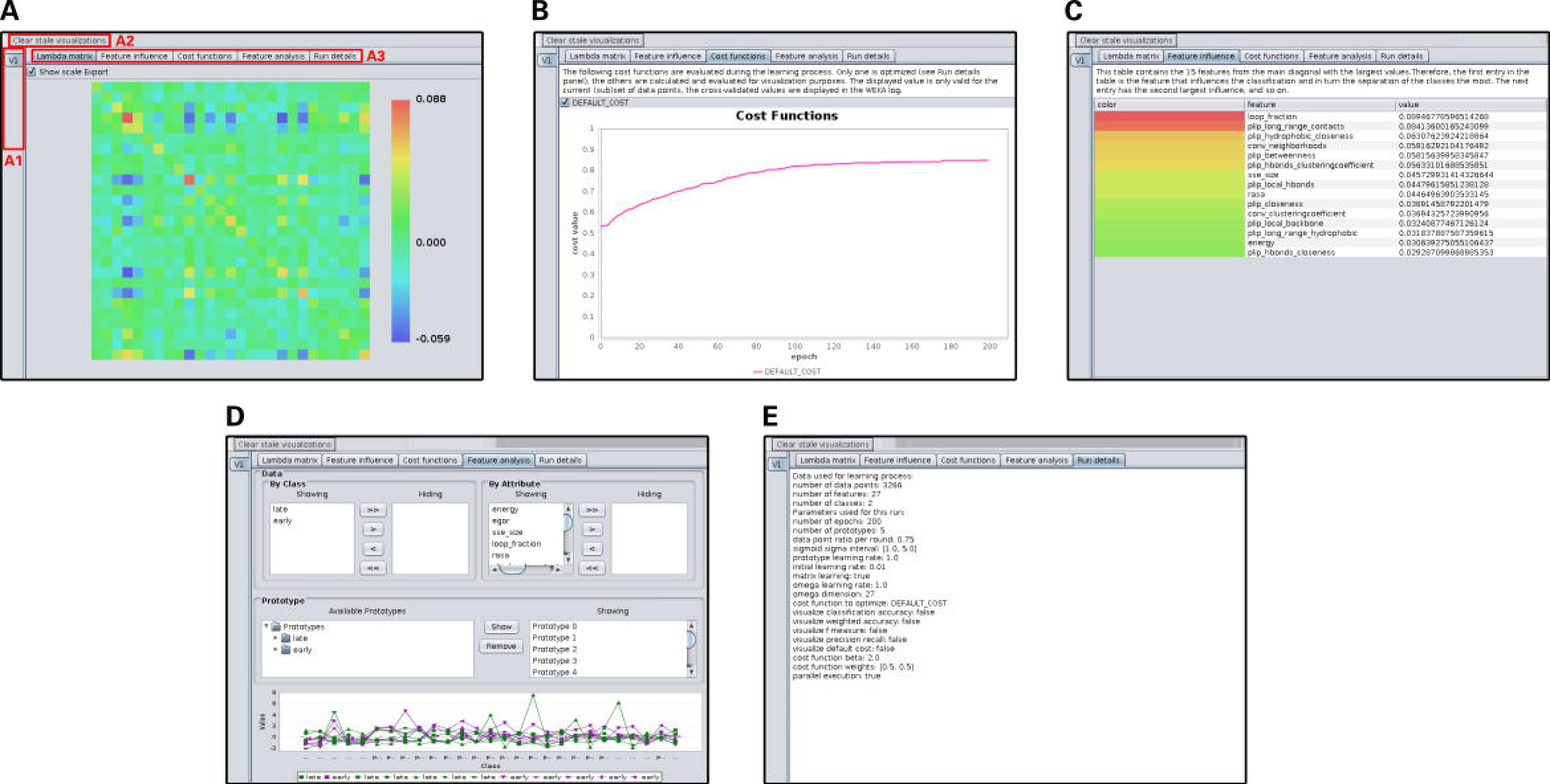
Screenshots of the Weka plug-in for GMLVQ. (A) The visualization of the CCM. The color scale indicates positive or negative correlations. (A1) The visualizations of each separate run will appear in this area. By clicking on the respective tab, one can easily switch between individual runs, e.g. cross validation runs. (A2) This button clears all visualizations except the latest. (A3) The tabs allow the user to switch between different visualizations of the current run. (B) The chart visualizes the cost functions over the course of learning. Additional functions can be visualized here, alongside with the cost function which is optimized. (C) The feature influence of single features of the current run. The top-ranked features have the highest contribution for the classification performance. (D) The feature analysis panel allows the detailed investigation of features and prototypes. (E) This panel shows the parameters which were used for the current run.

For the presented dataset, the CCM (Figure 5A) is primarily homogeneous which is indicated by values close to zero. The major contributing features are the *LF, PlipBN,* and especially *PlipHpCL* as these features exhibit the highest scores on the main diagonal of the CCM. The positive correlation of LF and *PlipBN* contributes to the classification performance as indicated by positive values described by the corresponding element. Also, the negative correlation of *PlipHpCL* to both features increases classification performance. The *PlipHpCL* is negatively correlated to various other features such as *SecSize, PlipLR, PlipHbLR,* and *PlipHbCL.* To a lesser degree, *e* and *PlipNC* are associated positively. It has to be pointed out that the CCM differs substantially from the correlation matrix (see Figure 5B). In the correlation matrix, strong positive correlations are present in the fourth group of features (local contact counts) and negative correlations in the fifth group (long-range contact counts). Relevant associations between features pointed out by GMLVQ are not obvious from the correlation matrix. The five most important features for discrimination are listed in Table 4 which was derived from the feature influence panel (Figure 6C). The prototype placement depicted in the feature analysis panel (Figure 6D) describes which values individual features adapt for optimal classification performance. This information is not evident from the CCM but necessary for the interpretation of the learned model. Selecting only these five features, and learning a model on this dimensionality-reduced dataset, shows a performance similar to the full model. GMLVQ with weighted accuracy and one prototype per class is given in Table 5. Recall and *F*_1_ value are even better compared to using all features. Thus, the GMLVQ can also used for feature extraction.

**Table 4.** Summary of the top five features, which are most important for the classification of EFR. The influence score is in arbitrary units, higher values refer to features which increase classification performance.

| feature | influence score |
| --- | --- |
| PlipHpCL | 0.159 |
| LF | 0.127 |
| PlipBN | 0.063 |
| SecSize | 0.059 |
| e | 0.042 |

**Table 5.** Performance using only the five most important features.

| CM |  | CA | PR | RE | F<sub>1</sub> | auROC |
| --- | --- | --- | --- | --- | --- | --- |
| 376 | 106 | 67.7 | 28.4 | 78.0 | 41.6 | 69.0 |
| 950 | 1834 |  |  |  |  |  |

The homogeneity observed in the CCM is the result of the similarity of several features. At a trivial level, topological descriptors computed on differing graph definitions are likely to result in redundant information. In that case, it is coincidental which feature will be highlighted even though all other correlated features capture similar information. Even if such features are strongly correlated, the CCM will only capture these characteristics if the correlation also contributes to the classification performance.

The *PlipBN* feature is the betweenness centrality [35, 36] derived from all contacts such as hydrogen bonds or hydrophobic interactions [33] in a protein structure. For this graph, residues with many of the shortest paths passing through them exhibit high betweenness centrality scores. This feature is highly discriminative for EFR and LFR as captured in the CCM. The prototypes which represent the EFR class display above average *PlipBN* values, indicating that EFR are better connected in the residue graph than their LFR counterparts. In fact, EFR exhibit a higher degree and are crucial connectors, so-called hubs. Residues with high betweenness centrality values have been shown to be crucial for the formation of stable, local structure and often constitute the folding nucleus of proteins [4, 36, 43].

The *LF* is relatively low for EFR which implies that EFR tend to be surrounded by ordered secondary structure elements. Analogously, this is negatively correlated to the size of the surrounding secondary structure elements and positively correlated to the *Rasa* values as it has been shown in previous studies [11, 12, 19, 36]. The *LF* feature is furthermore negatively correlated to *e* which indicates that ordered secondary structure elements result in favorable, low energy local conformations. These local structures are assumed to form autonomously and guide the folding process [12, 18].

The importance of the *PlipHpCL* represents the relevance of hydrophobic interactions in the core of protein structures (Figure 7). EFR have an increased propensity to occur in the core of protein structures which is isolated from the polar solvent [8, 19]. However, a buried or exposed state [44] derived from the *Rasa* feature cannot explain the origin and characteristics of EFR [17]. The closeness centrality [45] is defined as the inverse of the average path length of a residue to all other residues in the graph. It describes how well connected individual residues are which is a similar characteristic as covered by the betweenness centrality [35, 36]. The fact that both *PlipBN* and *PlipHpCL* are the most influential features for the classification demonstrates that they still capture slightly different aspects. The classification performance benefits from a negative correlation of both features. EFR occur primarily in the hydrophobic core of a structure where they participate in an increased number of hydrophobic interactions with surrounding residues. Previously, hydrophobic interactions have been shown to be relevant for the initiation and guidance of the protein folding process itself as well as its *in silico* modeling [2, 46–48]. They can be realized by a subset of amino acids and have an increased propensity to form ordered regions [11, 26]. The importance of the *PlipHpCL* feature and the placement of the prototypes implies that EFR are primarily embedded in the hydrophobic network of protein structures. EFR have been previously described to form more hydrophobic interactions which are important for the correct assembly of protein regions separated at sequence level [17].

**Figure 7.**
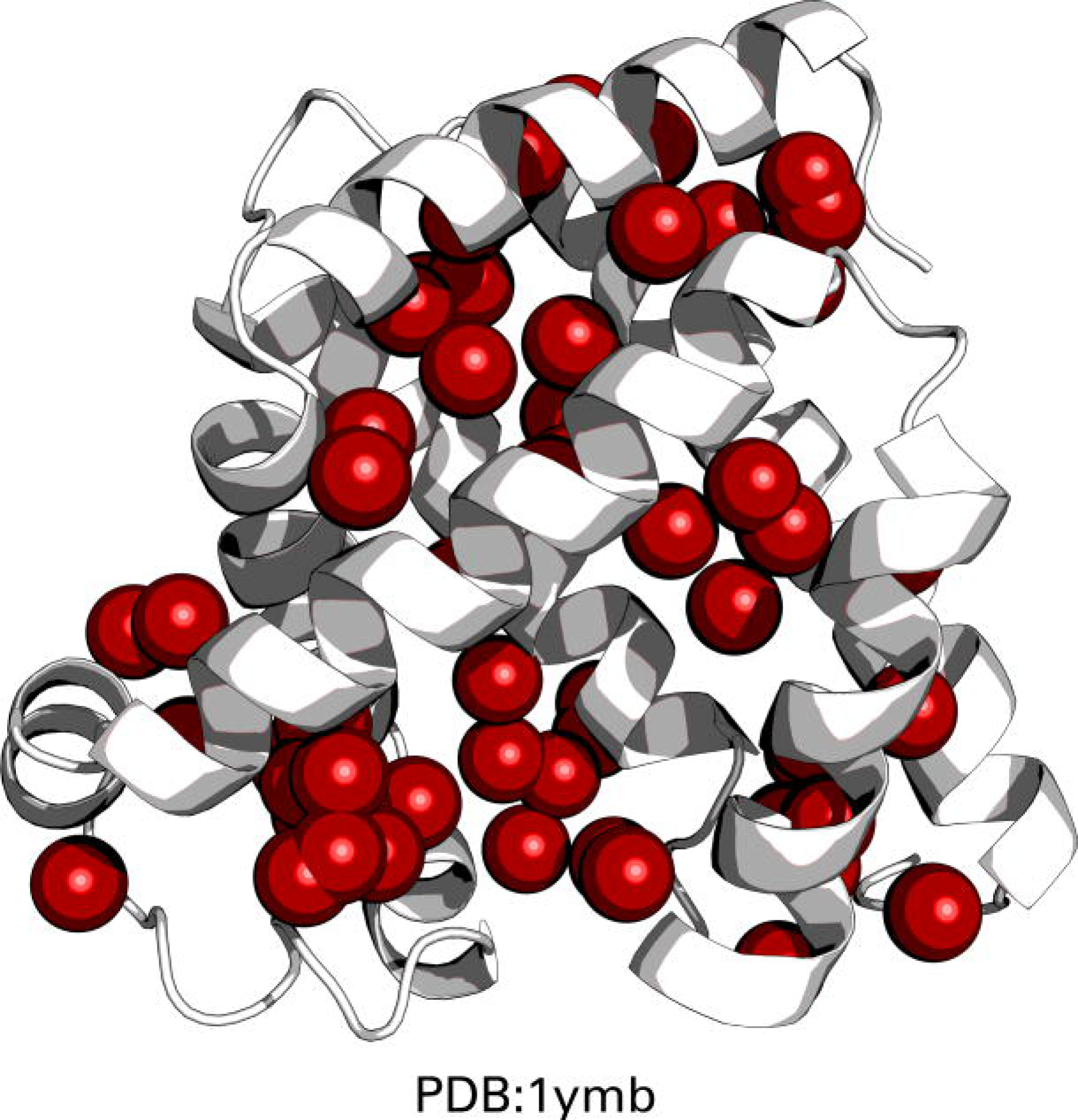
Rendering of the network of hydrophobic interactions. Structure of horse heart myoglobin (PDB:1ymb). In this structure, 58 hydrophobic interactions were detected by PLIP [33]. The centroids between interacting residues are depicted as red spheres. This highlights the strong contribution of hydrophobic interactions in the protein core.

In summary, the visualized classification of the GMLVQ run pointed out that many features capture redundant information. A subset of the features (*PlipHpCL, LF,* and *PlipBN*) is discriminative for both classes. Their importance and their respective correlations are in agreement with previous studies on EFR [16, 19] and, more general, folding nuclei [8, 12, 18, 36, 49].

## Conclusion

Machine as well as deep learning are trending in (life) sciences. Yet, a lot of classification problems are difficult to solve. Especially for problems with highly unbalanced class distributions the choice of the best model is crucial. Beside evaluation measures, other properties might be essential to select a suitable classifier. One key aspect is the interpretability of the learning process and the resulting model. GMLVQ is a prototype-based classifier. GMLVQ provides an interpretable classification model and was integrated into the Weka framework to make this classifier and its visualization capabilities accessible to a wide range of scientists.

A dataset of key residues of the protein folding process was investigated. GMLVQ performs on par with other state-of-the-art methods such as SVM or RF, but provides a readily interpretable classification model. From a set of 27 features, GMLVQ identified the fraction of ordered secondary structure elements, the betweenness centrality based on non-covalent contacts, and the closeness centrality using only hydrophobic interactions as the most relevant features for the distinction between Early and Late Folding Residues.

The classification performance may be improved by using additional features; however, for sake of simplicity such features were omitted because their computation would require additional algorithms or models. Promising candidates are backbone rigidity values [11], sequence based predictions of EFR [16], or evolutionary coupling scores [50]. All of them have been previously shown to be discriminative for Early Folding Residues [16, 19] and may increase the classification performance of this exemplary application of the Weka plug-in.

## Ethics approval and consent to participate

Not applicable.

## Consent for publication

Not applicable.

## Availability of data and material

All data generated or analyzed during this study is included in this published article and its supplementary information files. An open-source release and a step-by-step guide for the installation as well as usage of the GMLVQ Weka plug-in is available at https://github.com/JonStargaryen/gmlvq. The current version of the plug-in is deposited at https://doi.org/10.5281/zenodo.1326272.

## Competing interests

The authors declare that they have no competing interests.

## Funding

This work was funded by the European Social Fund (http://ec.europa.eu/esf/, grant numbers: 100235472 for FK and 1463643413126 for CL), the Free State of Saxony and the Saxon Ministry of Fine Arts. The funding agencies had no impact on design or conduct of the study.

## Author’s contributions

MK and TV designed the GMLVQ algorithm which was implemented in Java by SB, CL, and FK. SB and MK analyzed the data. SB, MK, CL, and FK drafted the manuscript. TV and DL supervised the project. All authors read and approved the manuscript.

## Acknowledgments

Not applicable.

**Tables**

**S1 File.**

Dataset in ARFF.

**S2 File.**

Dataset as CSV.

**S3 File.**

Correlation matrix of all features.

